# Osmosensing in trabecular meshwork cells

**DOI:** 10.1101/2024.04.03.587990

**Authors:** Jackson M. Baumann, Oleg Yarishkin, Monika Lakk, Christopher N. Rudzitis, Yun Ting Tseng, David Križaj

## Abstract

Aqueous humor drainage from the anterior eye constitutes a key determinant of intraocular pressure (IOP) under homeostatic and pathological conditions. Swelling of the trabecular meshwork (TM) increases its flow resistance but the mechanisms that sense and transduce osmotic gradients remain poorly understood. We used optical molecular analyses, optical imaging and electrophysiology to investigate TM osmotransduction and its role in calcium and chloride homeostasis. Anisosmotic conditions elicited proportional changes in TM cell volume. Swelling, but not shrinking, evoked increases in intracellular calcium concentration [Ca^2+^]_TM_. Hypotonicity-evoked calcium signals were sensitive to HC067047, a selective blocker of TRPV4 channels, whereas the agonist GSK1016790A promoted swelling under isotonic conditions. TRPV4 inhibition partially suppressed hypotonicity-induced volume increases and reduced the magnitude of the swelling-induced membrane current, with a substantial fraction of the swelling-evoked current abrogated by Cl^-^ channel antagonists DIDS and niflumic acid. The volume-sensing transcriptome of primary human TM cells showed expression of TRPV4, TRPM4, AQP1, and TMEMC3B genes. Cl^-^ channel expression was dominated by ANO6 transcripts, auxiliary levels of ANO3, ANO7 and ANO10 and modest expression of LTTRC genes that encode volume-activated anion channels. Thus, TRPV4-mediated cation influx works with Cl^-^ efflux to sense and respond to osmotic stress, potentially contributing to pathological swelling, calcium overload and intracellular signaling that could exacerbate functional disturbances in inflammatory disease and glaucoma.

## INTRODUCTION

The ability to sense and control intracellular volume represents one of the earliest and most highly conserved processes in cellular evolution. Cells swell in response to water influx driven by reduced ionic strength of the extracellular milieu, increased intracellular ion concentrations and/or upregulation of water-translocating transporters (Pasantes-Morales 2016, Toft-Bertelsen, Krizaj et al. 2017). Even small changes in intracellular water content and intra/extracellular volume influence protein function, diffusion of extracellular molecules and cellular performance (Epstein, Jedziniak et al. 1978, Pasantes-Morales 2016, Nicholson and Hrabetova 2017) while excessive swelling (edema) caused by ischemia, hyponatremia, traumatic brain/ocular/lung injury leads to energy depletion, loss of barrier function and oncotic, necrotic or apoptotic cell death (Platonova, Koltsova et al. 2012). Cytotoxic and vascular edema in the eye have been associated with pathologies such as corneal edema, diabetic retinopathy, Posner-Schlossman syndrome (PSS), age-related macular degeneration and vascular occlusion (Pannicke, Iandiev et al. 2006, Orduna Rios, Noguez Imm et al. 2019, Yan, Li et al. 2022, Martin-Gutierrez, Petzold et al. 2024). Healthy cells mitigate swelling-induced injury through “regulatory volume decrease” (RVD) - extrusion of Na^+^, K^+^, Cl^-^ ions and organic osmolytes that may be accompanied by changes in [Ca^2+^]_i_ (Hoffmann, Lambert et al. 2009).

Intraocular pressure (IOP) is homeostatically regulated by the trabecular meshwork (TM), a layered tissue composed of extracellular matrix (ECM) beams coated with contractile phagocytic cells that regulate the flow of aqueous fluid from the anterior chamber of the eye into the canal of Schlemm (SC) (Lutjen-Drecoll 1999, Stamer and Clark 2017). The TM resistance for the proto-glymphatic flow of aqueous humor is homeostatically adjusted by contractile activity of juxtacanalicular TM cells, which sense fluid osmolarity, shear and ECM strain (Ellingsen and Grant 1971, Krizaj 1995, Johnstone, Xin et al. 2021). The highly negative charges on sulfated glycosaminoglycan side chains of proteoglycans (aggrecan, versican) (Keller, Bradley et al. 2011) may generate osmotic pressure that facilitates the diffusion of aqueous humor across the TM. Given the quasi-linear inverse relationship between the size of the intracellular *vs.* extracellular space (Krizaj, Rice et al. 1996), any change in the volume of TM-resident cells will impact the hydraulic permeability of the trabecular pathway. The drainage of aqueous humor may indeed be affected by the concentration of soluble proteins and ions within the aqueous humor (Freddo, Patterson et al. 1984, Johnson, Gong et al. 1993, Gual, Llobet et al. 1997) and both mechanical loading and cell swelling may reduce the size of intercellular passages and impede conventional outflow in bovine, pig and human eyes (Gual, Llobet et al. 1997, Al-Aswad, Gong et al. 1999, Soto, Comes et al. 2004, Dismuke and Ellis 2009). Deficient volume regulation in glaucomatous TM cells suggests that sensing and management of osmotic gradients may be impaired in glaucoma (Gasull, Castany et al. 2019), and a water test has been developed to diagnose glaucoma risk in patients made to experience hypoosmotic shock (Susanna, Clement et al. 2017). Another example of the pathological relevance of TM swelling pertains to PSS patients, who may experience obstructed outflow due to trabeculitis and TM edema that can elevate IOP up to 40 mm Hg (Yan, Li et al. 2022). Among the many molecular mechanisms implicated in TM osmoresponsivity are Cl^-^, K^+^ and Na^+^ channels, NKCC transport, Na^+^/H^+^ antiport, nitric oxide synthase and aquaporins (AQPs) (Soto, Comes et al. 2004, Srinivas, Maertens et al. 2004, Comes, Abad et al. 2006, Baetz, Hoffman et al. 2009, Grant, Tran et al. 2013, Banerjee, Leung et al. 2017) but the identity and the signaling functions of the primary swelling sensor(s) remains poorly understood.

The objective of this study was to validate the significance of cell swelling for conventional outflow by characterizing novel volume sensing and transduction mechanisms in primary human TM cells that participate in [Ca^2+^]_TM_ homeostasis and swelling-induced ion transport. The presence of osmosensitive Ca^2+^ channels and/or transporters has been predicted by swelling- and Ca^2+^-activated K^+^ and Cl^-^ currents in bovine, pig and huma preparations (Mitchell, Fleischhauer et al. 2002, Soto, Comes et al. 2004, Srinivas, Maertens et al. 2004), but the source of the Ca^2+^ signal has remained unknown. We focused on TRPV4 (Transient Receptor Potential Vanilloid Isoform 4) due to its prominent expression in rodent and human TM (Luo, Conwell et al. 2014, Ryskamp, Frye et al. 2016), roles ins ensing pressure, strain and body temperature (Ryskamp, Frye et al. 2016, Yarishkin, Phuong et al. 2021, Krizaj, Cordeiro et al. 2023), role in TRPM4-mediated Na^+^ influx (Yarishkin, Phuong et al. 2022), AQP interactions (Jo, Ryskamp et al. 2015, Mola, Sparaneo et al. 2016) and sensitivity to hypotonic stimuli in other cell types (Liedtke and Friedman 2003, Hoffmann, Lambert et al. 2009). TRPV4 is a tetrameric channel with the S5-S6 region forming a pore permeable to cations, and functional domains within the N- and C-termini regulating trafficking, protein-protein interactions and sensitivity to lipid and mechanical milieus. Removal of the distal N-terminus results in loss of sensitivity to swelling whereas its replacement with a truncated domain of the cognate TRPV1 converted TRPV4 into a shrinking sensor (Toft-Bertelsen, Yarishkin et al. 2019). The channel has been implicated in osmosignaling in neurons, glia, epithelial cells, endothelial cells, heterologously expressing HEK293 cells and oocytes (Liedtke and Friedman 2003, Ryskamp, Jo et al. 2014, Phuong, Redmon et al. 2017, Toft-Bertelsen, Yarishkin et al. 2019, Lapajne, Lakk et al. 2020, Sonkusare and Laubach 2022) but not keratinocytes (Ritzmann, Jahn et al. 2023). TRPV4 modulates systemic osmoregulation (Liedtke and Friedman 2003), water diffusion and edema formation in the retina, lung and brain (Hoshi, Okabe et al. 2018, Orduna Rios, Noguez Imm et al. 2019, Sonkusare and Laubach 2022) but its role in volume regulation has been under debate, with studies implicating it in the promotion of swelling (Jo, Ryskamp et al. 2015, Barile, Mola et al. 2023) and regulatory volume decrease (RVD) (Becker, Blase et al. 2005, Hoffmann, Lambert et al. 2009, Jo, Ryskamp et al. 2015). Here, we identify TRPV4 as a central transducer of swelling-induced calcium signaling in human TM cells, show that its activation contributes to volume expansion, and elucidate its collaboration with Cl^-^ efflux as a component of the swelling-evoked current I_Swell_.

## MATERIALS & METHODS

### TM cell culture

De-identified postmortem eyes from 5 male and female donors aged 60 - 81 years with no history of glaucoma were procured from Utah Lions Eye Bank with written informed consent of the donors’ families and according to the standards set by the WMA Declaration of Helsinki and the Department of Health and Human Services Belmont Report. Cells from juxtacanalicular and corneoscleral TM regions were isolated (Yarishkin, Phuong et al. 2018, Lakk and Krizaj 2021, Yarishkin, Phuong et al. 2022) and cultured, showing the typical spindle and hexagonal phenotypes (Stamer and Clark 2017). The cells were validated following the consensus recommendations (Keller, Bhattacharya et al. 2018), including periodic profiling for MYOC, AQP1, MGP, ACTA2 expression and dexamethasone-dependent upregulation of MYOC (Lakk and Krizaj 2021, Yarishkin, Phuong et al. 2022). Passage 2–6 cells were seeded onto collagen I coverslips and grown in Trabecular Meshwork Cell Medium (ScienCell Research Laboratories, Carlsbad, CA) supplemented with 2% fetal bovine serum (FBS), 100 U/ml penicillin, 100 mg/ml streptomycin, at pH 7.4, 37°C and 5% CO_2_.

### Animals

Studies were conducted using enucleated eyes from C57BL6 mice. The animals were euthanized in accordance with the NIH Guide for the Care and Use of Laboratory Animals, the ARVO Statement for the Use of Animals in Ophthalmic and Vision Research, and the Institutional Animal Care and Use Committees at the University of Utah and Duke University (IACUC approvals 19-04005 and A184-18-08, respectively). The animals were maintained in pathogen-free facilities with a 12-hour light/dark cycle, *ad libitum* access to food and water and temperature set to ∼22-23°C. Because mice do not exhibit sex differences in facility, data from male and female animals was pooled.

### Reagents

Most salts and reagents, including chloride channel blockers DIDS (4,4’-Diisothiocyanato-2,2’-stilbenedisulfonic acid) and niflumic acid were purchased from Sigma-Aldrich (St. Louis, MO, USA). GSK1016790A (GSK101) and HC067047 (HC-06) were obtained from Cayman Biotech. *Grammostola spatulata* mechanotoxin 4 (GsMTx4) was from Alomone Labs Aliquots of GSK101 (1-10 mM) and HC-06 (4 mM) stocks in DMSO were diluted 1:1000 in extracellular saline before use.

### PCR analysis

Total RNA was isolated using the Arcturus RNeasy Plus Micro Kit (Qiagen) (Lakk et al., 2021). 1µg of total RNA was used for reverse transcription. First-strand cDNA synthesis and PCR amplification of cDNA were performed using qScript ^TM^ Ultra SuperMix cDNA synthesis kit (Quanta Biosciences). SYBR Green based real-time PCR was performed using Apex qPCR Master Mix (Genesee Scientific). PCR conditions were as follows: 95°C for 15 min, 95°C for 20 s, 60°C for 60 s, 40 cycles. The results were performed in triplicate of at least four three experiments. The comparative C _T_ method (ΔΔC_T_) was used to measure relative gene expression where the fold enrichment was calculated as: 2 − ^[*Δ*CT^ ^(sample)−^ *^Δ^*^CT^ ^(calibrator)]^ after normalization. *Gapdh* was utilized to normalize fluorescence signals. The primer sequences and expected product sizes are given in Table I.

### Optical imaging

TM cells were seeded onto glass coverslips for 48 h, loaded with fluorescent dyes (Calcein AM or Fura-2 AM; ThermoFisher Scientific) for 40–60 min, and washed with the isotonic bath solution containing (in mM): 57.5 NaCl, 153 mannitol, 2.5 KCl, 1.2 MgCl_2_, 5.6 glucose, 10 HEPES, 1.8 CaCl_2_ (pH 7.4, osmolarity 300 mOsm). Hypotonic solutions were prepared by removing requisite amounts of mannitol from the extracellular solution (15 and 110 mM, respectively) and the hypertonic solution by addition of 100 mM mannitol (Ryskamp, Jo et al. 2014, Jo, Ryskamp et al. 2015). Saline was delivered to the recording chamber through a gravity-fed 8-reservoir system (Warner Instruments/Harvard Bioscience, Holliston, MA). Osmolarity was checked with a vapor pressure osmometer (Wescor, Logan, UT).

Cell volume was measured as fluorescence in cells loaded with 10 mM Calcein AM (ThermoFisher Scientific) at RT for 30 min. The dye was excited at 488 nm with emission at 520 nm tracked with an EMCCD camera (Delta Evolve, Photometrics, Phoenix, AZ) and NIS Elements software (Nikon Instruments, Redmon, WA). Every (isotonic, anisotonic) experimental condition was assessed with 3-5 slides/experiment and 10 – 15 cells/slide (i.e. 90 – 255 cells/condition) with the data representing results averaged across at least 3 experiments.

For calcium imaging, cells were loaded with 5-10 µM of Fura-2 AM with trapping by de-esterification assumed to accumulate the intracellular concentration to ∼100 µM (Krizaj and Copenhagen 1998). Epifluorescence images were acquired on an inverted Nikon Ti microscope with 20x (0.75 N.A. oil) and 40x (1.3 N.A. oil & 0.8 N.A. water) objectives. Excitation for 340 nm and 380 nm filters (Semrock, Rochester, NY, USA) was delivered by a liquid light guide from a 150 W xenon arc lamp (DG-4, Sutter Instruments, Novato, CA, USA). Fluorescence emission was high pass-filtered at 510 nm and captured with a cooled digital CCD camera (Photometrics Delta, Tucson, AZ). Ca^2+^ signals were plotted as ΔR/R (peak F_340_/F_380_ ratio – baseline/baseline), with *F*_340_/*F*_380_ calculations and background subtraction performed on Regions of Interest (ROI) that encompassed the central cell area (NIS Elements 3.22 or 5.0; Nikon Instruments) (Ryskamp, Frye et al. 2016, Lakk, Young et al. 2018, Lapajne, Lakk et al. 2020, Redmon, Yarishkin et al. 2021). Experiments were performed at room temperature (20–22°C).

### Electrophysiology

Borosilicate patch pipettes (WPI) were pulled using a P-2000 micropipette puller (Sutter Instruments, Novato, CA) from borosilicate glass (1.5 mm O.D., 0.84 mm I.D.) with electrode resistance of 6–8 MΩ when filled with the internal buffer solution containing (mM): 125 K-gluconate, 10 KCl, 1.5 MgCl_2_, 10 HEPES, 10 ethylene glycol-bis(β-aminoethyl ether)-N,N, N’,N”-tetraacetic acid (EGTA), pH 7.4. The chamber was superfused with saline containing (in mM): 140 NaCl, 2.5 KCl, 1.5 MgCl_2_, 1.5 CaCl_2_, 5.6 D-glucose, 10 HEPES (pH 7.4, adjusted with NaOH) (Yarishkin, Phuong et al. 2018, Redmon, Yarishkin et al. 2021). The holding potential in the whole-cell recordings was set to −40 mV. The membrane capacitance of TM cells was measured with pClamp 10.6 software (Molecular Devices, San Jose, CA).

Whole-cell currents were acquired with a Multiclamp 700B amplifier, pClamp 10.6 software and Digidata 1440A interface (Molecular Devices). Data were sampled at 5 kH, digitized at 2 kH, and analyzed with Clampfit 10.7 (Molecular Devices) and Origin 8 Pro (OriginLab, Northampton, MA). The patch clamp experiments were performed at room temperature (20–22°C).

### Data analysis

All data are presented as the means ± S.EM. Student’s paired *t-test*, two-sample *t* test or ANOVA multiple comparisons test were applied to estimate statistical significance of results. *P* < 0.05 was considered statistically significant. Some of the data in this paper have been presented in an abstract form (Baumann et al., 2019).

## RESULTS

### TRPV4 regulates hypotonicity-evoked TM swelling

Cellular volume balances passive distribution of water with active mechanisms that maintain the osmotic steady state (Hoffmann, Lambert et al. 2009, Pasantes-Morales 2016). Calcein-AM quenching was used in combination with ratiometric imaging to assess the response of human TM cells to a physiological (285 mOsm) and pathological hypotonic (190 mOsm) gradients and a hypertonic (400 mOsm) stimulus. Anisosmotic stimuli evoked dose-dependent changes in calcein fluorescence: 285 mOsm perfusates increased the volume signal by 6.7 ± 1.1% (n = 141; P < 0.01), 190 mOsm perfusates by 23.2 ± 3.4% (n = 134; P < 0.005) and hypertonic 400 mOsm saline induced −19.8 ± 5.1% decrease in fluorescence (n = 118; P < 0.005) (Fig. 1). The amplitude of HTS-evoked fluorescence was reproducible, with sustained (10 min) exposure to hypotonic gradients not associated with observable RVD under our experimental conditions (Fig. 1A). An unexpected side effect of pathological swelling were membrane blebs that appeared across the cell body, occasionally detached, and disappeared following the return to isotonicity (Fig. 1C).

**Figure 1.**
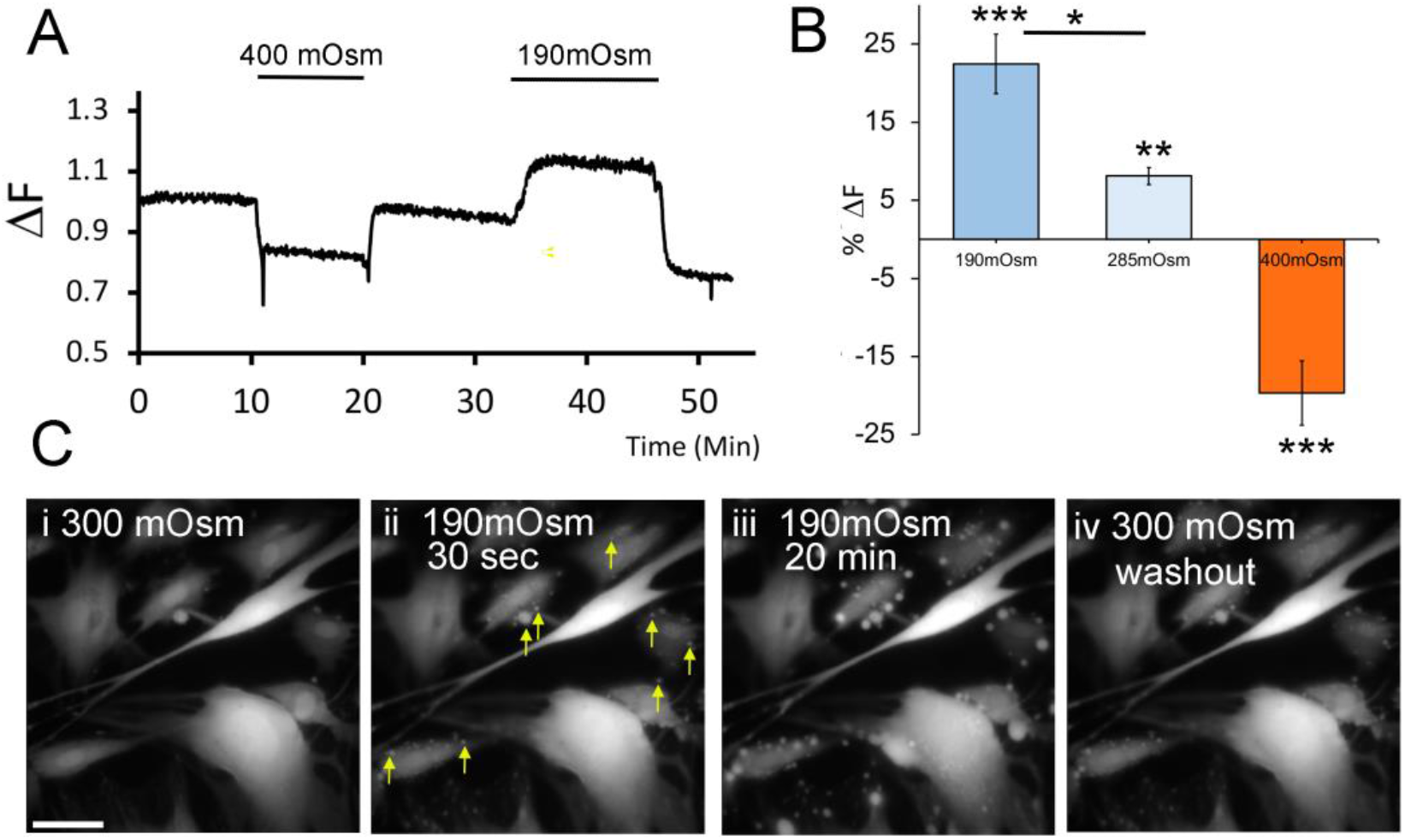
TM cell volume as an osmometer: anisotonic conditions alter the cell volume. A Fluorescence traces from calcein AM-loaded pTM cells exposed to hypotonic (190 mOsm) and hypertonic (400 mOsm) saline. B Averaged fluorescence data for hypotonicity (285 mOsm, 190 mOsm) and hypertonicity (400 mOsm)-induced changes in fluorescence. C Time series of TM cell response to 190 mOsm. Blebs appear within seconds of HTS imposition (arrows), are maintained during the imposed stimulus and wash out following the return to isotonic saline. (N = 3, n = 118 – 134).

Studies of volume- and Ca^2+^-sensitive anion channels and BK (KCa1.1) K^+^ channels in TM cells (Soto, Comes et al. 2004, Banerjee, Leung et al. 2017) predicted TM swelling to be associated with cytosolic influx of Ca^2+^. Given the expression of TRPV4 (Luo, Conwell et al. 2014, Ryskamp, Frye et al. 2016), its functions as a swelling sensor (Liedtke and Friedman 2003, Iuso and Krizaj 2016, Toft-Bertelsen, Krizaj et al. 2017) and transducer of TM strain (Ryskamp, Frye et al. 2016, Yarishkin, Phuong et al. 2021), we hypothesized it also participates in TM volume sensing and regulation. Figure 2D shows that the selective TRPV4 antagonist HC067047 (HC-06, 5 μM) antagonizes swelling induced by hypotonic stimulation (HTS), with the extent of cell volume increase reduced from 23.2% ± 3.4% to 14.2% ± 1.3% in the presence of HC-06 (P < 0.05). The selective agonist GSK1016790A (GSK101, 25 nM) applied under isotonic conditions, decreased calcein-AM fluorescence by 25.4% ± 6.3% (P = 0.05) (Fig. 2B). The effect of isotonic GSK101 superfusion on TM cell volume was comparable to the effect of 190 mOsm saline.

**Figure 2.**
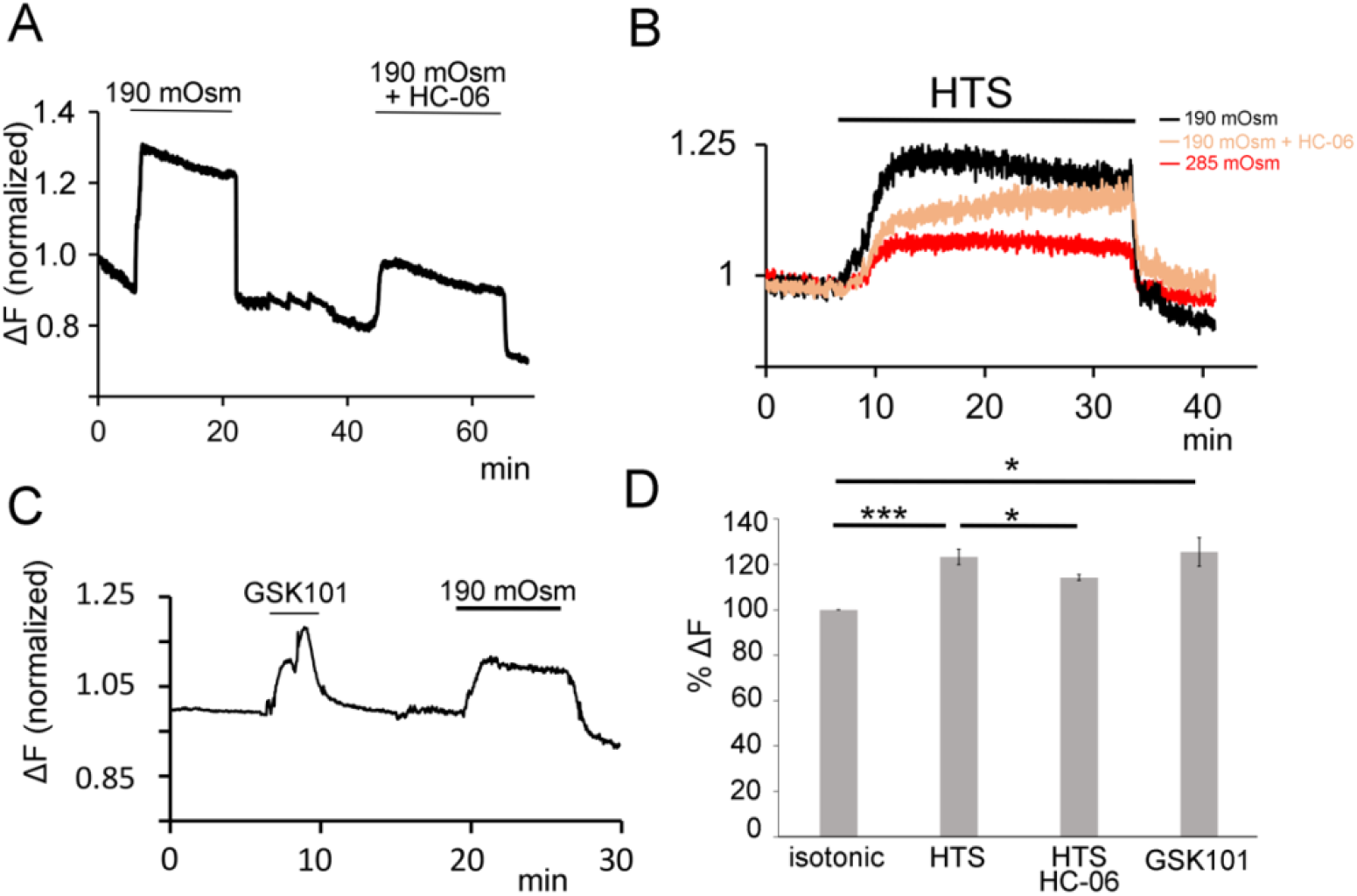
Hypotonicity-induced swelling has a TRPV4 Component. Calcein AM-loaded cells were stimulated with HTS in the presence/absence of TRPV4 channel modulators. A & B The amplitude of HTS (190 mOsm) -evoked swelling is reduced in the presence of the TRPV4 antagonist HC-06.C Representative example of calcein fluorescence. TRPV4 agonist GSK101 (25 nM) and HTS (190 mOsm) induced comparable increases in pTM cell volume. D Averaged data for the experimental conditions in A-C. HTS (190 mOsm)-induced increases in cell volume are reduced in the presence of HC-06, and superfusion with GSK101 is associated with cell swelling under isotonic conditions (N = 3, n = 72 – 165). * P < 0.05, *** P < 0.005

### TRPV4 mediates the response to physiological osmolarity gradients and regulates RVD

A nonselective cation channel (P_Ca_/P_Na_ ∼6-10) (Liedtke and Friedman 2003, Deng, Paknejad et al. 2018), TRPV4 mediates Ca^2+^ influx upon activation. We tracked this process in pTM cells loaded with the ratiometric indicator Fura-2 AM (5 μM) and superfused with anisotonic saline. TM cells responded to HTS with [Ca^2+^]i elevations that were TRPV4-dependent, as indicated by the reduced amplitude in the presence of HC-06 (Fig. 4). The TRPV4 antagonist suppressed the amplitude of the ratiometric signal from 0.93 ± 0.11 (n = 115) to 0.28 ± 0.07 (n = 101; P < 0.01), ∼70% inhibition. In contrast, hypertonic stimulation was not accompanied by changes in [Ca^2+^]_i_ (N = 3, data not shown).

### Transduction of cell swelling involves collaboration between cationic mechanochannels

Pressure-induced cation influx in TM cells consists of time-dependent activation of TRPV41 and Piezo1 currents (Yarishkin, Phuong et al. 2021). Generally associated with strain transduction, Piezo1 has been shown to mediate sensitivity to swelling in heterologously expressing HEK293 (Sforna, Michelucci et al. 2022). We tested Piezo1 involvement in HTS-induced signaling by exposing TM cells to 190 mOsm saline in the presence of HC-06 concentrations that block TRPV4 channels (5 μM) (Yarishkin, Phuong et al. 2021)) and GsMTx4, a tarantula toxin antagonist of Piezo1 (Gnanasambandam, Gottlieb et al. 2017, Yarishkin, Phuong et al. 2021). GsMTx4 (5 μM) induced a modest but significant (P = 0.048) decrease in the amplitude of [Ca^2+^]_i_ signals (from 0.28 ± 0.07 to 0.19 ± 0.1) (n = 71) (Fig. 3D).

**Figure 3.**
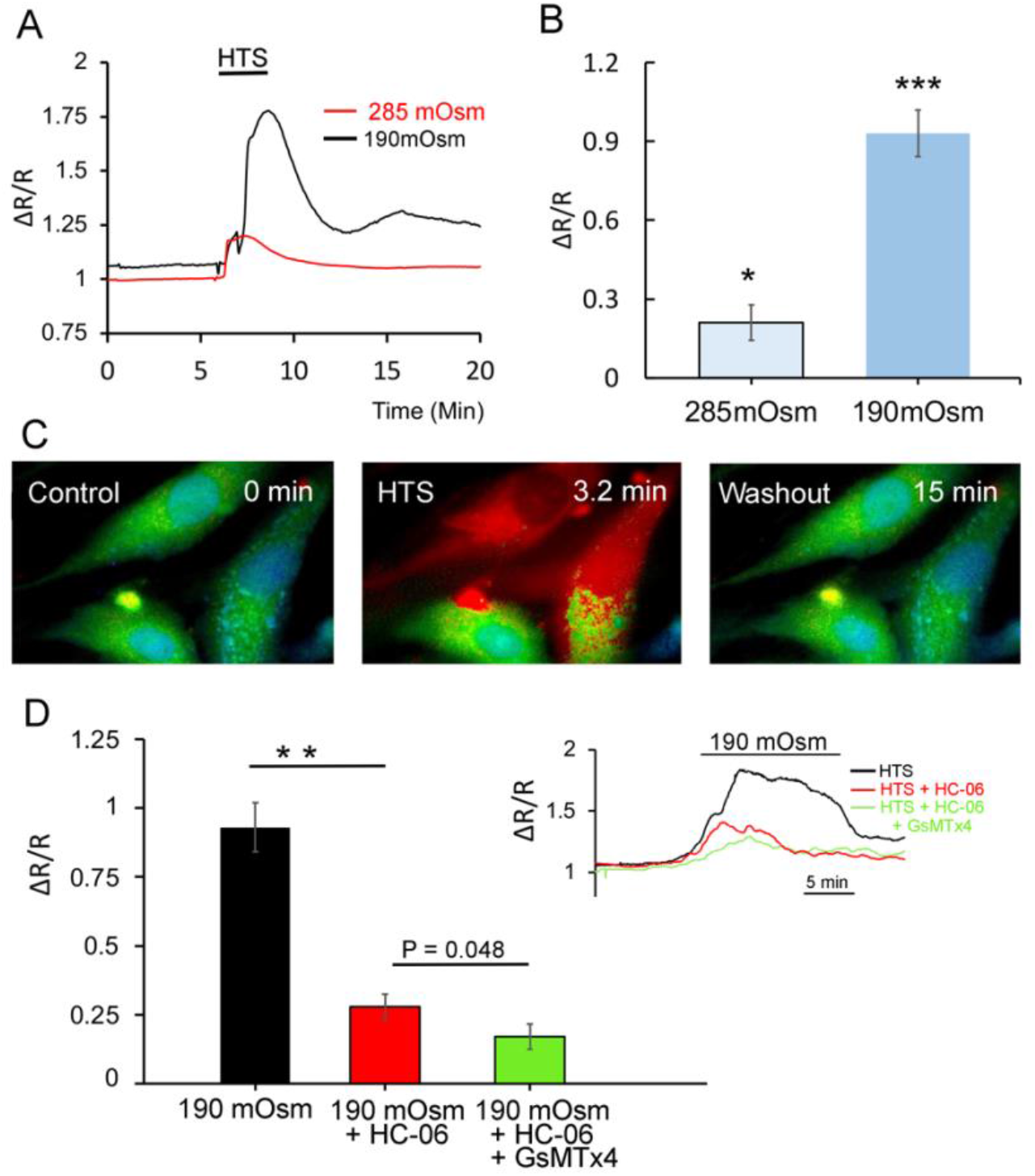
Hypotonicity evokes [Ca^2+^]_i_ elevations that are partially mediated by TRPV4. A Representative time course of HTS-evoked in [Ca^2+^]_TM_, with the amplitude of the [Ca^2+^]_i_ response reflecting the magnitude of the imposed gradient. B Averaged data for the experiments in A. C Spatial representation of Fura-2 fluorescence before (left panel), during (middle panel) and following (right panel) cell exposure to 190 mOsm saline. D Averaged data from 3 independent experiments, 71-115 cells/experimental condition. HTS (190 mOsm)-evoked [Ca^2+^]_i_ response was significantly reduced by the TRPV4 antagonist HC-06, with GsMTx4 producing a additional reduction in response amplitude (N = 3, 71-115 cells). Inset: time course of HTS-evoked ratiometric signals under control conditions (black trace), during superfusion with HC-06 and combined superfusion with HC-06 + GsMTx4.

### TRPV4 and chloride channels together mediate I_Swell_

Calcium signals mediated by TRPV4 activation in TM cells include transmembrane and Ca^2+^-Induced Ca^2+^ release (CICR) components (Ryskamp, Frye et al. 2016). To characterize TRPV4-mediated membrane influx we measured the HTS-evoked current in cells under whole-cell voltage-clamp. HTS evoked a substantial (nA!) swelling-evoked current (I_Swell_) with reversal at ∼-25 mV. The time course of I_Swell_ induced by 190 mOsm is shown in Figure 4A at the holding potentials of ±100 mV.

At the resting potential (−40 mV (Yarishkin, Phuong et al. 2018, Yarishkin, Phuong et al. 2022)) I_Swell_ was inward and slow, taking min to reach the peak amplitude of –329 pA ± 45 pA (N =18). 285 mOsm saline evoked currents with correspondingly reduced amplitude. Pretreatment with HC-06 attenuated the amplitude of I_Swell_ by ∼ 36.2% ± 8% (N = 10; P < 0.05) (Fig. 4F), indicating that a substantial component of I_TMSwell_ is mediated by TRPV4.

**Figure 4.**
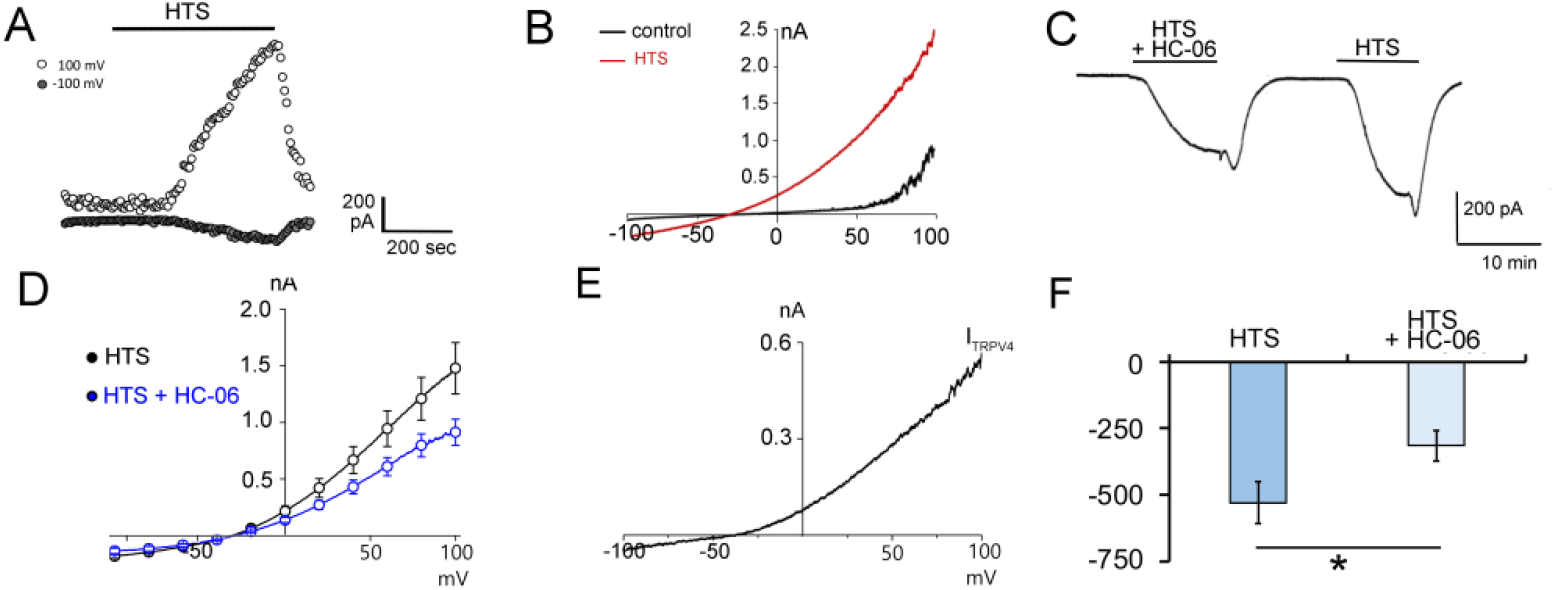
TRPV4 partially contributes to I_Swell_. Whole cell recording, voltage clamp. A Mean amplitude time course of inward and outward currents induced by HTS −100 mV (white circles) and +100 mV (grey circles). B Voltage-dependence of I_Swell_ (red trace) shows modest outward rectification. C Representative time course of HTS-evoked current shows slow onset and suppression by HC-06. D I-V relationship of the HTS-evoked current in the presence and absence of HC-06 shows modest suppression by HC-06. E The I-V relationship of HTS-activated I_TRPV4_ (i.e., HC-06-sensitive current). F Averaged I_Swell_ data. The amplitude of the evoked current was partially reduced by HC-06 (P < 0.05; n = 10).

Previous studies of TM osmosignaling generally associated it with chloride signaling and transport (Al-Aswad, Gong et al. 1999, Mitchell, Fleischhauer et al. 2002, Soto, Comes et al. 2004, Srinivas, Maertens et al. 2004, Banerjee, Leung et al. 2017, Gasull, Castany et al. 2019). Consistent with this, the nonselective Cl^-^ channel blocker DIDS (300 μM) reduced the amplitude of the HTS-evoked current by ∼70% (n = 5) (Fig. 6A). Applied in conjunction with HC-06, DIDS abrogated the HTS-evoked current (P < 0.01; n = 9). Coapplication of HC-06 and niflumic acid similarly eliminated I_Swell_ (Fig. 5C & D). I_Swell_ thus consists of Na^+^ and Ca^2+^ influx mediated by TRPV4 channels and Cl- flux mediated by volume-activated Cl^-^ channels.

**Figure 5.**
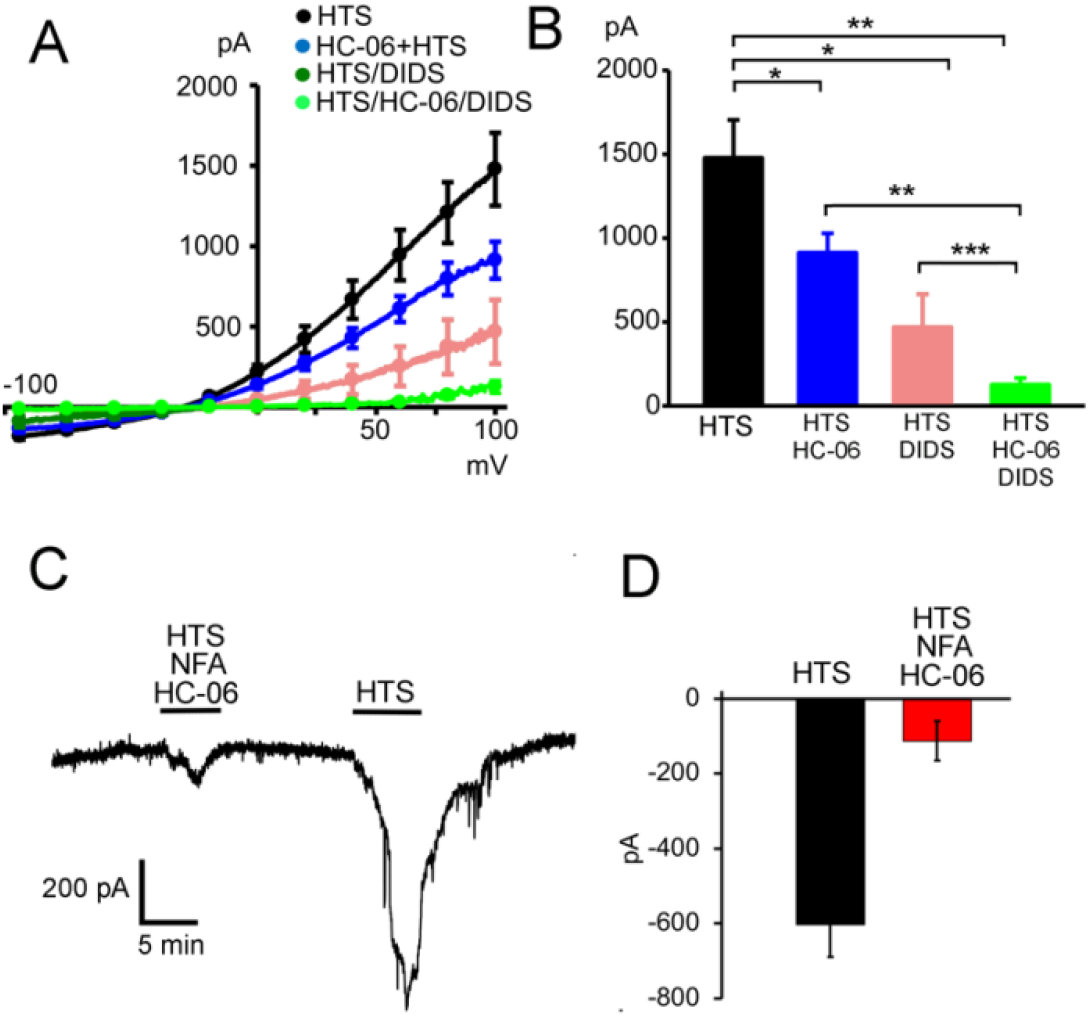
I_Swell_ is mediated by TRPV4 and Cl- channels. A & B I-V relationship and peak current amplitude for I_Swell_ under indicated conditions (HTS, 190 mOsm). I_Swell_ is inhibited by combined application of HC-06 and DIDS. C & D HTS stimulation does not evoke a current in the presence of HC- 06 + niflumic acid (NFA). * P < 0.05, ** P < 0.01, *** P < 0.005 (N = 3, n = 5 – 9/condition)

**Figure 6.**
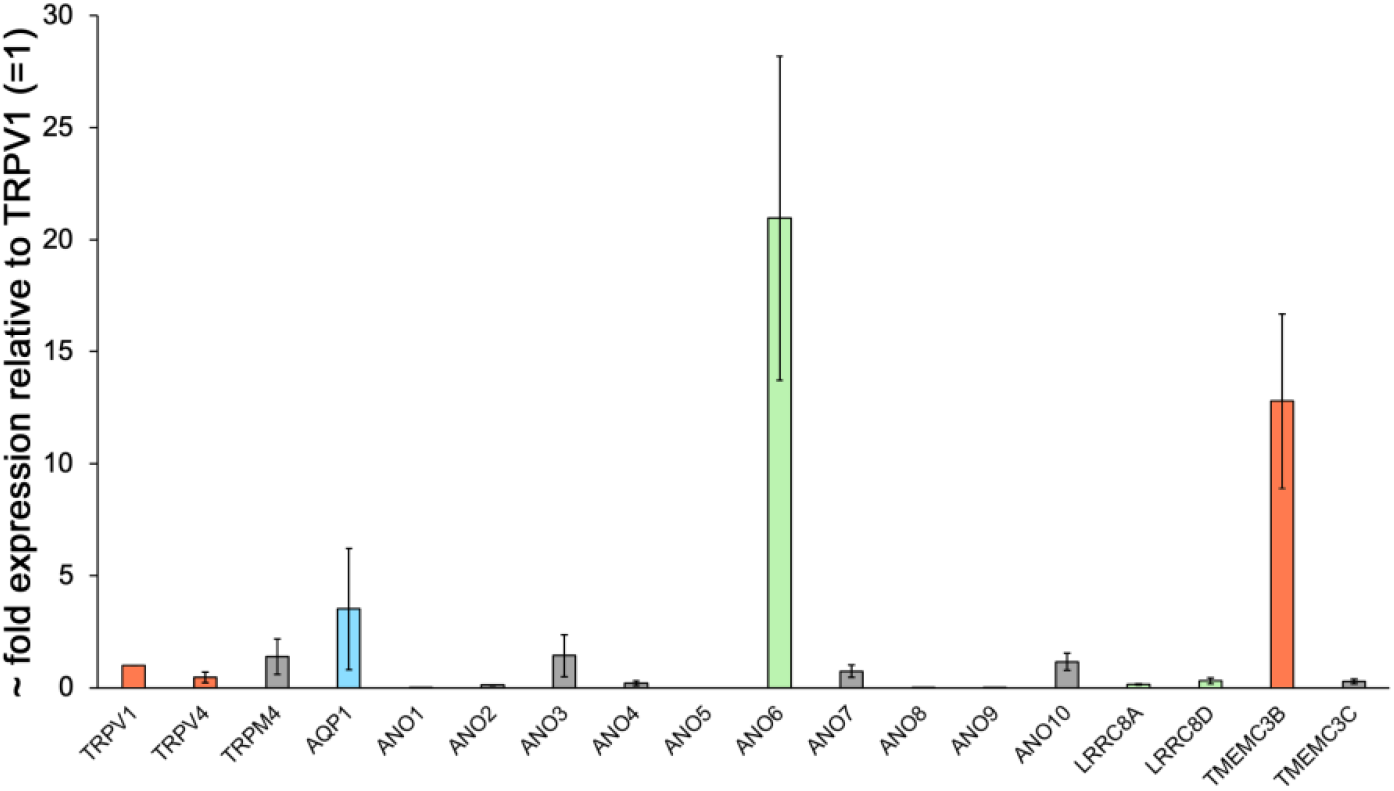
Relative expression levels of genes that encode swelling-relevant proteins. pTM cells show prominent expression ANO6, residual expression of ANO3, ANO7 and ANO10, and modest expression of LRRC8 subunits. Expression of TMEM63B transcripts is prominent (N = 3). Data normalized vs. TRPV1 levels.

### Expression of Osmo-relevant genes in human TM cells

Previous studies associated TM swelling with activation of VRAC and anoctamin channels (Soto, Comes et al. 2004, Banerjee, Leung et al. 2017, Gasull, Castany et al. 2019). We used semiquantitative qPCR to gain insight into relative expression of anoctamins (TMEM16) and VRAC subunit genes, TRPM4 and AQP1 genes, as well as the potential expression of TMEM63B transcripts that encode a novel osmoactivated Ca^2+^ channel (Du, Ye et al. 2020) homologous to plant OSCA channels (Murthy, Dubin et al. 2018) with no known function in the eye. As expected, pTM cells showed TRPV4, TRPM4 and AQP1 expression. ANO6 mRNAs dominated anoctamin transcript levels alongside the auxiliary expression of anoctamins 3, 7, and 10. pTM cells prominently expressed TMEM63B, but levels of LRTTC8A/8D mRNAs were lower compared to ANO6 transcripts. These data point at intricate multi-channel Ca^2+^ and Cl^-^ volume signaling in mammalian TM cells.

## DISCUSSION

The results reported here indicate that sensing and transduction of osmotic stress within the TM requires activation of TRPV4 channels, which mediate swelling-induced [Ca^2+^]_i_ signals and collaborate with volume-sensitive Cl^-^ efflux to carry the swelling-induced transmembrane current. Our findings bring insight into the mechanisms that mediate TM osmosensitivity in the presence of physiological and pathological gradients, and could therefore participate in the modulation of outflow in mammalian eyes.

We found that TM cells function as osmometers that, as reported for many plant, invertebrate and vertebrate preparations (Krizaj, Rice et al. 1996, Hoffmann, Lambert et al. 2009, Pasantes-Morales 2016, Nicholson and Hrabetova 2017), track imposed ionic gradients with proportional changes in cell volume (Fig. 2) (Mitchell, Fleischhauer et al. 2002). pTM cells utilize TRPV4, a nonselective cation channel previously associated with IOP regulation (Luo, Conwell et al. 2014, Ryskamp, Frye et al. 2016) as the principal conduit for swelling-induced influx of Ca^2+^ and thus the likely source of osmosensitive Ca^2+^- activated K^+^ and Cl- fluxes predicted by electrophysiological studies (Mitchell, Fleischhauer et al. 2002, Soto, Comes et al. 2004, Banerjee, Leung et al. 2017). In addition to mediating hypotonicity-induced increases in free [Ca^2+^]_i_, TRPV4 also collaborates with Cl^-^ channels to carry I_Swell_. TM cell shrinking was not associated with [Ca^2+^]_i_ changes, suggesting that TRPV1 channels are not represented by the truncated hypertonicity-sensitive N-terminal variant (Prager-Khoutorsky and Bourque 2015, Toft-Bertelsen, Yarishkin et al. 2019). The observations that TRPV4 inhibition suppresses HTS-induced increases in [Ca^2+^]_TM_, reduces the extent of cell swelling (Fig. 2 & 3) and reduces the amplitude of I_Swell_ (Fig. 5) identify TRPV4 as a principal transducer of osmotic swelling in mammalian TM cells. Swelling-induced TRPV4 activation has been suggested to require the distal N-terminus of the protein, cytoskeletal tethering and/or upstream activation of phospholipase A2 (PLA2) that culminates in production of epoxyeicosatrienoic acids (EETs) as final common activators of the channel (Watanabe, Vriens et al. 2003, Vriens, Watanabe et al. 2004, Hoffmann, Lambert et al. 2009, Toft-Bertelsen, Yarishkin et al. 2019). While HTS-induced TRPV4 activation appears to be independent of PLA2, actin and phosphorylation in some cells (Ryskamp, Jo et al. 2014, Toft-Bertelsen, Krizaj et al. 2017, Toft-Bertelsen, Yarishkin et al. 2019) and the channel may not require membrane tethering and biosynthesis of polyunsaturated fatty acids (Ryskamp, Frye et al. 2016), the slow onset of the HTS-evoked current (Fig. 4 & 5) and pressure-evoked current (Yarishkin, Phuong et al. 2021) are consistent with time-dependent formation of EET messenger molecules (Vriens, Watanabe et al. 2004, Ryskamp, Frye et al. 2016).

TRPV4-mediated osmosensing has been associated with the promotion of cell swelling and RVD (Becker, Blase et al. 2005, Hoffmann, Lambert et al. 2009, Jo, Ryskamp et al. 2015, Barile, Mola et al. 2023). The suppression of the amplitude of HTS-induced swelling by HC-06 and stimulation of cell volume expansion by GSK101 under isotonic conditions suggest that, as reported recently for glial cells (Jo, Ryskamp et al. 2015, Iuso and Krizaj 2016, Mola, Sparaneo et al. 2016, Barile, Mola et al. 2023) TRPV4 activity promotes cell swelling instead of time-dependent volume regulation. The increase in osmotic permeability during TRPV4 activation could reflect Ca^2+^-dependent stimulation of TRPM4 channels, NKCC transporters, release of polyunsaturated fatty acids and/or an increase in the rate of cell swelling due to AQP-mediated water transport. We and others found that coexpression of TRPV4 and AQP1 facilitates cell swelling, whereas ablation of AQP1/AQP4 channels reduces the rate and extent of swelling in transfected oocytes and glia (Jo, Ryskamp et al. 2015, Mola, Sparaneo et al. 2016, Toft-Bertelsen, Krizaj et al. 2017). Given that (i) TRPV4 inhibition, knockdown and Ca^2+^ chelation suppress HTS-induced swelling (Fig. 2) (Jo, Ryskamp et al. 2015) and (ii) swelling is augmented in cells that overexpress TRPV4 (Barile, Mola et al. 2023), AQPs may not be required to facilitate cell swelling in response TRPV4-mediated Ca^2+^ influx. NKCC1 transports water (Zeuthen and MacAulay 2002) and may modulate outflow resistance (Al-Aswad, Gong et al. 1999) whereas TRPM4 shows strong expression in mouse and human TM and mediates 9-phenanthrol-sensitive Na^+^ influx in GSK101-stimulated pTM cells (Yarishkin, Phuong et al. 2022). Cell swelling may be inhibited by TRPM4 ablation and inhibition (Stokum, Kwon et al. 2018). Finally, activation of TM TRPV4 channels stimulates the release of arachidonic acid (Uchida, Shimizu et al. 2021), a TRPV4 agonist (Ryskamp, Frye et al. 2016) that may promote cell swelling through a positive feedback loop (Ryskamp, Iuso et al. 2015). TRPV4 inhibitors alleviate cytotoxic edema in the brain, lung and retina (Hoshi, Okabe et al. 2018, Orduna Rios, Noguez Imm et al. 2019, Sucha, Hermanova et al. 2022, Tureckova, Hermanova et al. 2023) but could also be utilized to mitigate TM swelling during inflammatory conditions that compromise the facility (e.g., glaucomatocyclitic edema (Yan, Li et al. 2022)). Minimal RVD under our experimental conditions (Fig. 2) argues against a major role for Ca^2+^-dependent efflux of ions and organic osmolytes (Becker, Blase et al. 2005).

Given that HTS-evoked [Ca^2+^]_i_ signals in TM cells were not completely antagonized by HC-06 concentrations that block GSK101-evoked Ca^2+^ responses (Ryskamp, Frye et al. 2016, Lakk and Krizaj 2021, Yarishkin, Phuong et al. 2021), the swelling response may involve additional Ca^2+^ influx pathways - such as Piezo1 channels, which mediate the response to shear flow in TM cells (Yarishkin, Phuong et al. 2021) and swelling-induced Ca^2+^ signaling in heterologously expressing cells (Sforna, Michelucci et al. 2022). Combined inhibition of TRPV4 and Piezo1 channels was associated with a modest [Ca^2+^]_i_ decrease, with the residual (∼10 - 25%) signal suggesting potential activation of yet-to-be characterized plasma membrane or Ca^2+^ store mechanisms. We note that pTM cells express TMEM63B transcripts encoding a novel Ca^2+^-permeable channel that has been associated with transduction of hypotonic (Du, Ye et al. 2020) and hypertonic (Yang, Jia et al. 2024) stimuli in animals and plants. The absence of Ca^2+^ elevations in hypertonicity-treated cells argues against its role in cell shrinking but we cannot exclude its potential contribution to the swelling-induced calcium signaling.

We found that I_TMSwell_ can be abrogated by co-application of HC-06 and niflumic acid/DIDS. DIDS and NFA do not affect the TRPV4 current (Rahman, Mukherjee et al. 2016), indicating that the HC-06- and DIDS-sensitive components are mediated by mediating nonselective cation influx via TRPV4 (∼35% of I_Swell_) and Cl^-^ efflux (∼65% of I_Swell_) via yet to be identified Cl^-^ channels. Cells often express multiple classes of Cl^-^ channels and transporters, which regulate the membrane potential, development and signaling and may include an osmosensitive subset (Duran, Thompson et al. 2010). The sensitivity of I_Swell_ to DIDS and its mild outward rectification accord with the properties of VRAC (Akita and Okada 2014) which has been proposed to mediate swelling-induced Cl^-^ flux in mammalian TM cells (Mitchell, Fleischhauer et al. 2002, Gasull, Castany et al. 2019). However, the expression of LTTRC8A/LTRRCE subunits was surprisingly modest and the minimal RVD and weak voltage inactivation of I_TMSwell_ at positive potentials (Fig. 4A) argue against major VRAC contributions. The transcriptomic profile of pTM cells was dominated by ANO6 (TMEM16F) transcripts, which encode Ca^2+^-activated Cl^-^ channels that also function as phospholipid scramblases (Duran, Thompson et al. 2010). In trophoblasts, GSK101 activates the ANO6 current together with phospholipid scrambling (Zhang, Liang et al. 2022), suggesting the possibility of cation and anion flux interactions. Consistent with the role for ANO6 in mediating the HTS-induced current, its knockdown resulted in 50-70% I_Swell_ reduction of in immortalized TM cells (Banerjee, Leung et al. 2017). The delineation of chloride transport mechanisms during TM signaling under normal and pathological conditions will need to consider ANO6, VRAC heteromers, together with ANO3, ANO7, and ANO10 proteins that are transcribed in pTM cells, may function as scramblases (Nguyen and Chen 2024) and participate in intracellular Cl^-^ transport (Duran, Thompson et al. 2010) yet remain to be studied in the eye. TM cells may express voltage-gated Cl^-^ channels such as ClC2 (Comes, Abad et al. 2006) but their poor antagonism for DIDS (Alexander and Grinstein 2006) and lack of I_Swell_ inward rectification argue against significant contribution to HTS-induced Cl^-^ efflux.

Another notable aspect of HTS-associated cellular signaling in TM cells was the reversible formation of blebs that occasionally detached from the plasma membrane and disappeared during isotonic washout. Similar vacuole-like membrane reservoirs have been described in fibroblasts that experienced reductions in extracellular tonicity (Zhang, Gao et al. 2000, Charras 2008, Beyder and Sachs 2009, Kosmalska, Casares et al. 2015), and were proposed to reflect the breakdown of cortical F-actin and β-tubulin (Kosmalska, Casares et al. 2015). Blebs were proposed to be enriched with caveolins and mechanosensitive channels and thus could show increased sensitivity to tensile stress as the cell membrane expands in response to ionic flux (Cox, Bae et al. 2016). pTM membranes contain ∼4 distinct TRPV4 peptides, with the 3 large- M.W. variants excluded from caveolae and lipid rafts (Lakk, Hoffmann et al. 2021). However, the truncated ∼75kDa TRPV4 variant co-immunoprecipitates with Cav- 1 (Lakk, Hoffmann et al. 2021), suggesting the possibility that the TM membrane partitions into areas with differential mechanosensitivities. Whether HTS-sensitive blebs in TM cells correspond to actin-free and/or caveolar domains (e.g.,(Charras 2008, Kosmalska, Casares et al. 2015, Cox, Bae et al. 2016)) remains to be seen in future work.

In summary, this study reports that TRPV4 channels mediate the principal calcium signal fraction in swollen TM cells, contribute to I_Swell_ and that their activity promotes swelling and presumably contributes to suppression of aqueous fluid drainage. Volume as an extensive thermodynamic parameter is presumably never directly sensed by a cell, which instead responds to its changes by tracking the tensile status of the cell membrane with stretch-sensitive channels such as TRPV4. TRPV4 activity could contribute to impaired volume regulation in glaucoma (Susanna, Clement et al. 2017), trabeculitic edema (Yan, Li et al. 2022) and symptoms of renal failure, which is accompanied by osmotic imbalances between the serum and aqueous humor (Kilavuzoglu, Yurteri et al. 2018). Targeting TRPV4 channels could potentially alleviate TM edema under inflammatory conditions (Yan, Li et al. 2022). Future studies may investigate the precise role of chloride efflux and transport and assess the biological functions of SLC transporters that comprise over 400 proteins, many of which with minimally defined functions. Finally, as a polymodal transducer of pressure, ECM strain, polyunsaturated fatty acids, cholesterol, and temperature (Ryskamp, Frye et al. 2016, Lakk, Hoffmann et al. 2021, Yarishkin, Phuong et al. 2021) TRPV4 participates in multiple aspects of sensory transduction within the TM, acting as an interface between mechanosensory, volume signaling, fatty acid and contractile mechanisms.

## GRANTS

Supported by the National Institutes of Health (R01EY027920, R01EY031817, P30EY014800, T32EY024234 to D.K.; T32EY024234 to C.N.R), Crandall Glaucoma Initiative, Stauss-Rankin Foundation, and unrestricted support from Research to Prevent Blindness to Moran Eye Institute at the University of Utah and Duke University.

## Notes

### Competing Interest Statement

The authors have declared no competing interest.

## REFERENCES

Akita, T. and Y. Okada (2014). “Characteristics and roles of the volume-sensitive outwardly rectifying (VSOR) anion channel in the central nervous system.” Neuroscience 275: 211–231.

Al-Aswad, L. A., H. Gong, D. Lee, M. E. O’Donnell, J. D. Brandt, W. J. Ryan, A. Schroeder and K. A. Erickson (1999). “Effects of Na-K-2Cl cotransport regulators on outflow facility in calf and human eyes in vitro.” Invest Ophthalmol Vis Sci 40(8): 1695–1701.

Alexander, R. T. and S. Grinstein (2006). “Na+/H+ exchangers and the regulation of volume.” Acta Physiol (Oxf) 187(1-2): 159–167.

Baetz, N. W., E. A. Hoffman, A. J. Yool and W. D. Stamer (2009). “Role of aquaporin-1 in trabecular meshwork cell homeostasis during mechanical strain.” Exp Eye Res 89(1): 95–100.

Banerjee, J., C. T. Leung, A. Li, K. Peterson-Yantorno, H. Ouyang, W. D. Stamer and M. M. Civan (2017). “Regulatory Roles of Anoctamin-6 in Human Trabecular Meshwork Cells.” Invest Ophthalmol Vis Sci 58(1): 492–501.

Barile, B., M. G. Mola, F. Formaggio, E. Saracino, A. Cibelli, C. D. Gargano, G. Mogni, A. Frigeri, M. Caprini, V. Benfenati and G. P. Nicchia (2023). “AQP4-independent TRPV4 modulation of plasma membrane water permeability.” Front Cell Neurosci 17: 1247761.

Becker, D., C. Blase, J. Bereiter-Hahn and M. Jendrach (2005). “TRPV4 exhibits a functional role in cell-volume regulation.” J Cell Sci 118(Pt 11): 2435–2440.

Beyder, A. and F. Sachs (2009). “Electromechanical coupling in the membranes of Shaker-transfected HEK cells.” Proc Natl Acad Sci U S A 106(16): 6626–6631.

Charras, G. T. (2008). “A short history of blebbing.” J Microsc 231(3): 466–478.

Comes, N., E. Abad, M. Morales, T. Borras, A. Gual and X. Gasull (2006). “Identification and functional characterization of ClC-2 chloride channels in trabecular meshwork cells.” Exp Eye Res 83(4): 877–889.

Cox, C. D., C. Bae, L. Ziegler, S. Hartley, V. Nikolova-Krstevski, P. R. Rohde, C. A. Ng, F. Sachs, P. A. Gottlieb and B. Martinac (2016). “Removal of the mechanoprotective influence of the cytoskeleton reveals PIEZO1 is gated by bilayer tension.” Nat Commun 7: 10366.

Deng, Z., N. Paknejad, G. Maksaev, M. Sala-Rabanal, C. G. Nichols, R. K. Hite and P. Yuan (2018). “Cryo-EM and X-ray structures of TRPV4 reveal insight into ion permeation and gating mechanisms.” Nat Struct Mol Biol 25(3): 252–260.

Dismuke, W. M. and D. Z. Ellis (2009). “Activation of the BK(Ca) channel increases outflow facility and decreases trabecular meshwork cell volume.” J Ocul Pharmacol Ther 25(4): 309–314.

Du, H., C. Ye, D. Wu, Y. Y. Zang, L. Zhang, C. Chen, X. Y. He, J. J. Yang, P. Hu, Z. Xu, G. Wan and Y. S. Shi (2020). “The Cation Channel TMEM63B Is an Osmosensor Required for Hearing.” Cell Rep 31(5): 107596.

Duran, C., C. H. Thompson, Q. Xiao and H. C. Hartzell (2010). “Chloride channels: often enigmatic, rarely predictable.” Annu Rev Physiol 72: 95–121.

Ellingsen, B. A. and W. M. Grant (1971). “The relationship of pressure and aqueous outflow in enucleated human eyes.” Invest Ophthalmol 10(6): 430–437.

Epstein, D. L., J. A. Jedziniak and W. M. Grant (1978). “Obstruction of aqueous outflow by lens particles and by heavy-molecular-weight soluble lens proteins.” Invest Ophthalmol Vis Sci 17(3): 272–277.

Freddo, T. F., M. M. Patterson, D. R. Scott and D. L. Epstein (1984). “Influence of mercurial sulfhydryl agents on aqueous outflow pathways in enucleated eyes.” Invest Ophthalmol Vis Sci 25(3): 278–285.

Gasull, X., M. Castany, A. Castellanos, M. Rezola, A. Andres-Bilbe, M. I. Canut, R. Estevez, T. Borras and N. Comes (2019). “The LRRC8-mediated volume-regulated anion channel is altered in glaucoma.” Sci Rep 9(1): 5392.

Gnanasambandam, R., P. A. Gottlieb and F. Sachs (2017). “The Kinetics and the Permeation Properties of Piezo Channels.” Curr Top Membr 79: 275–307.

Grant, J., V. Tran, S. K. Bhattacharya and L. Bianchi (2013). “Ionic currents of human trabecular meshwork cells from control and glaucoma subjects.” J Membr Biol 246(2): 167–175.

Gual, A., A. Llobet, R. Gilabert, M. Borras, J. Pales, M. V. Bergamini and C. Belmonte (1997). “Effects of time of storage, albumin, and osmolality changes on outflow facility (C) of bovine anterior segment in vitro.” Invest Ophthalmol Vis Sci 38(10): 2165–2171.

Hoffmann, E. K., I. H. Lambert and S. F. Pedersen (2009). “Physiology of cell volume regulation in vertebrates.” Physiol Rev 89(1): 193–277.

Hoshi, Y., K. Okabe, K. Shibasaki, T. Funatsu, N. Matsuki, Y. Ikegaya and R. Koyama (2018). “Ischemic Brain Injury Leads to Brain Edema via Hyperthermia-Induced TRPV4 Activation.” J Neurosci 38(25): 5700–5709.

Iuso, A. and D. Krizaj (2016). “TRPV4-AQP4 interactions ‘turbocharge’ astroglial sensitivity to small osmotic gradients.” Channels (Austin) 10(3): 172–174.

Jo, A. O., D. A. Ryskamp, T. T. Phuong, A. S. Verkman, O. Yarishkin, N. MacAulay and D. Krizaj (2015). “TRPV4 and AQP4 Channels Synergistically Regulate Cell Volume and Calcium Homeostasis in Retinal Muller Glia.” J Neurosci 35(39): 13525–13537.

Johnson, M., H. Gong, T. F. Freddo, N. Ritter and R. Kamm (1993). “Serum proteins and aqueous outflow resistance in bovine eyes.” Invest Ophthalmol Vis Sci 34(13): 3549–3557.

Johnstone, M., C. Xin, J. Tan, E. Martin, J. Wen and R. K. Wang (2021). “Aqueous outflow regulation - 21st century concepts.” Prog Retin Eye Res 83: 100917.

Keller, K. E., S. K. Bhattacharya, T. Borras, T. M. Brunner, S. Chansangpetch, A. F. Clark, W. M. Dismuke, Y. Du, M. H. Elliott, C. R. Ethier, J. A. Faralli, T. F. Freddo, R. Fuchshofer, M. Giovingo, H. Gong, P. Gonzalez, A. Huang, M. A. Johnstone, P. L. Kaufman, M. J. Kelley, P. A. Knepper, C. C. Kopczynski, J. G. Kuchtey, R. W. Kuchtey, M. H. Kuehn, R. L. Lieberman, S. C. Lin, P. Liton, Y. Liu, E. Lutjen-Drecoll, W. Mao, M. Masis-Solano, F. McDonnell, C. M. McDowell, D. R. Overby, P. P. Pattabiraman, V. K. Raghunathan, P. V. Rao, D. J. Rhee, U. R. Chowdhury, P. Russell, J. R. Samples, D. Schwartz, E. B. Stubbs, E. R. Tamm, J. C. Tan, C. B. Toris, K. Y. Torrejon, J. A. Vranka, M. K. Wirtz, T. Yorio, J. Zhang, G. S. Zode, M. P. Fautsch, D. M. Peters, T. S. Acott and W. D. Stamer (2018). “Consensus recommendations for trabecular meshwork cell isolation, characterization and culture.” Exp Eye Res 171: 164–173.

Keller, K. E., J. M. Bradley, J. A. Vranka and T. S. Acott (2011). “Segmental versican expression in the trabecular meshwork and involvement in outflow facility.” Invest Ophthalmol Vis Sci 52(8): 5049–5057.

Kilavuzoglu, A. E. B., G. Yurteri, N. Guven, S. Marsap, A. R. C. Celebi and C. B. Cosar (2018). “The effect of hemodialysis on intraocular pressure.” Adv Clin Exp Med 27(1): 105–110.

Kosmalska, A. J., L. Casares, A. Elosegui-Artola, J. J. Thottacherry, R. Moreno-Vicente, V. Gonzalez-Tarrago, M. A. Del Pozo, S. Mayor, M. Arroyo, D. Navajas, X. Trepat, N. C. Gauthier and P. Roca-Cusachs (2015). “Physical principles of membrane remodelling during cell mechanoadaptation.” Nat Commun 6: 7292.

Krizaj, D. (1995). What is glaucoma? Webvision: The Organization of the Retina and Visual System. H. Kolb, E. Fernandez and R. Nelson. Salt Lake City (UT).

Krizaj, D. and D. R. Copenhagen (1998). “Compartmentalization of calcium extrusion mechanisms in the outer and inner segments of photoreceptors.” Neuron 21(1): 249–256.

Krizaj, D., S. Cordeiro and O. Strauss (2023). “Retinal TRP channels: Cell-type-specific regulators of retinal homeostasis and multimodal integration.” Prog Retin Eye Res 92: 101114.

Krizaj, D., M. E. Rice, R. A. Wardle and C. Nicholson (1996). “Water compartmentalization and extracellular tortuosity after osmotic changes in cerebellum of Trachemys scripta.” J Physiol 492 (Pt 3)(Pt 3): 887–896.

Lakk, M., G. F. Hoffmann, A. Gorusupudi, E. Enyong, A. Lin, P. S. Bernstein, T. Toft-Bertelsen, N. MacAulay, M. H. Elliott and D. Krizaj (2021). “Membrane cholesterol regulates TRPV4 function, cytoskeletal expression, and the cellular response to tension.” J Lipid Res 62: 100145.

Lakk, M. and D. Krizaj (2021). “TRPV4-Rho signaling drives cytoskeletal and focal adhesion remodeling in trabecular meshwork cells.” Am J Physiol Cell Physiol 320(6): C1013–C1030.

Lakk, M., D. Young, J. M. Baumann, A. O. Jo, H. Hu and D. Krizaj (2018). “Polymodal TRPV1 and TRPV4 Sensors Colocalize but Do Not Functionally Interact in a Subpopulation of Mouse Retinal Ganglion Cells.” Front Cell Neurosci 12: 353.

Lapajne, L., M. Lakk, O. Yarishkin, L. Gubeljak, M. Hawlina and D. Krizaj (2020). “Polymodal Sensory Transduction in Mouse Corneal Epithelial Cells.” Invest Ophthalmol Vis Sci 61(4): 2.

Liedtke, W. and J. M. Friedman (2003). “Abnormal osmotic regulation in trpv4-/- mice.” Proc Natl Acad Sci U S A 100(23): 13698–13703.

Luo, N., M. D. Conwell, X. Chen, C. I. Kettenhofen, C. J. Westlake, L. B. Cantor, C. D. Wells, R. N. Weinreb, T. W. Corson, D. F. Spandau, K. M. Joos, C. Iomini, A. G. Obukhov and Y. Sun (2014). “Primary cilia signaling mediates intraocular pressure sensation.” Proc Natl Acad Sci U S A 111(35): 12871–12876.

Lutjen-Drecoll, E. (1999). “Functional morphology of the trabecular meshwork in primate eyes.” Prog Retin Eye Res 18(1): 91–119.

Martin-Gutierrez, M. P., A. Petzold and Z. Saihan (2024). “NAION or not NAION? A literature review of pathogenesis and differential diagnosis of anterior ischaemic optic neuropathies.” Eye (Lond) 38(3): 418–425.

Mitchell, C. H., J. C. Fleischhauer, W. D. Stamer, K. Peterson-Yantorno and M. M. Civan (2002). “Human trabecular meshwork cell volume regulation.” Am J Physiol Cell Physiol 283(1): C315–326.

Mola, M. G., A. Sparaneo, C. D. Gargano, D. C. Spray, M. Svelto, A. Frigeri, E. Scemes and G. P. Nicchia (2016). “The speed of swelling kinetics modulates cell volume regulation and calcium signaling in astrocytes: A different point of view on the role of aquaporins.” Glia 64(1): 139–154.

Murthy, S. E., A. E. Dubin, T. Whitwam, S. Jojoa-Cruz, S. M. Cahalan, S. A. R. Mousavi, A. B. Ward and A. Patapoutian (2018). “OSCA/TMEM63 are an Evolutionarily Conserved Family of Mechanically Activated Ion Channels.” Elife 7.

Nguyen, D. M. and T. Y. Chen (2024). “Structure and Function of Calcium-Activated Chloride Channels and Phospholipid Scramblases in the TMEM16 Family.” Handb Exp Pharmacol 283: 153–180.

Nicholson, C. and S. Hrabetova (2017). “Brain Extracellular Space: The Final Frontier of Neuroscience.” Biophys J 113(10): 2133–2142.

Orduna Rios, M., R. Noguez Imm, N. M. Hernandez Godinez, A. M. Bautista Cortes, D. D. Lopez Escalante, W. Liedtke, A. Martinez Torres, L. Concha and S. Thebault (2019). “TRPV4 inhibition prevents increased water diffusion and blood-retina barrier breakdown in the retina of streptozotocin-induced diabetic mice.” PLoS One 14(5): e0212158.

Pannicke, T., I. Iandiev, A. Wurm, O. Uckermann, F. vom Hagen, A. Reichenbach, P. Wiedemann, H. P. Hammes and A. Bringmann (2006). “Diabetes alters osmotic swelling characteristics and membrane conductance of glial cells in rat retina.” Diabetes 55(3): 633–639.

Pasantes-Morales, H. (2016). “Channels and Volume Changes in the Life and Death of the Cell.” Mol Pharmacol 90(3): 358–370.

Phuong, T. T. T., S. N. Redmon, O. Yarishkin, J. M. Winter, D. Y. Li and D. Krizaj (2017). “Calcium influx through TRPV4 channels modulates the adherens contacts between retinal microvascular endothelial cells.” J Physiol 595(22): 6869–6885.

Platonova, A., S. V. Koltsova, P. Hamet, R. Grygorczyk and S. N. Orlov (2012). “Swelling rather than shrinkage precedes apoptosis in serum-deprived vascular smooth muscle cells.” Apoptosis 17(5): 429–438.

Prager-Khoutorsky, M. and C. W. Bourque (2015). “Mechanical basis of osmosensory transduction in magnocellular neurosecretory neurones of the rat supraoptic nucleus.” J Neuroendocrinol 27(6): 507–515.

Rahman, M., S. Mukherjee, W. Sheng, B. Nilius and L. J. Janssen (2016). “Electrophysiological characterization of voltage-dependent calcium currents and TRPV4 currents in human pulmonary fibroblasts.” Am J Physiol Lung Cell Mol Physiol 310(7): L603–614.

Redmon, S. N., O. Yarishkin, M. Lakk, A. Jo, E. Mustafic, P. Tvrdik and D. Krizaj (2021). “TRPV4 channels mediate the mechanoresponse in retinal microglia.” Glia 69(6): 1563–1582.

Ritzmann, D., M. Jahn, S. Heck, C. Jung, T. Cesetti, N. Couturier, R. Rudolf, N. Reuscher, C. Buerger, O. Rauh and T. Fauth (2023). “The Ca(2+) channel TRPV4 is dispensable for Ca(2+) influx and cell volume regulation during hypotonic stress response in human keratinocyte cell lines.” Cell Calcium 111: 102715.

Ryskamp, D. A., A. M. Frye, T. T. Phuong, O. Yarishkin, A. O. Jo, Y. Xu, M. Lakk, A. Iuso, S. N. Redmon, B. Ambati, G. Hageman, G. D. Prestwich, K. Y. Torrejon and D. Krizaj (2016). “TRPV4 regulates calcium homeostasis, cytoskeletal remodeling, conventional outflow and intraocular pressure in the mammalian eye.” Sci Rep 6: 30583.

Ryskamp, D. A., A. Iuso and D. Krizaj (2015). “TRPV4 links inflammatory signaling and neuroglial swelling.” Channels (Austin) 9(2): 70–72.

Ryskamp, D. A., A. O. Jo, A. M. Frye, F. Vazquez-Chona, N. MacAulay, W. B. Thoreson and D. Krizaj (2014). “Swelling and eicosanoid metabolites differentially gate TRPV4 channels in retinal neurons and glia.” J Neurosci 34(47): 15689–15700.

Sforna, L., A. Michelucci, F. Morena, C. Argentati, F. Franciolini, M. Vassalli, S. Martino and L. Catacuzzeno (2022). “Piezo1 controls cell volume and migration by modulating swelling-activated chloride current through Ca(2+) influx.” J Cell Physiol 237(3): 1857–1870.

Sonkusare, S. K. and V. E. Laubach (2022). “Endothelial TRPV4 channels in lung edema and injury.” Curr Top Membr 89: 43–62.

Soto, D., N. Comes, E. Ferrer, M. Morales, A. Escalada, J. Pales, C. Solsona, A. Gual and X. Gasull (2004). “Modulation of aqueous humor outflow by ionic mechanisms involved in trabecular meshwork cell volume regulation.” Invest Ophthalmol Vis Sci 45(10): 3650–3661.

Srinivas, S. P., C. Maertens, L. H. Goon, L. Goon, M. Satpathy, B. Y. Yue, G. Droogmans and B. Nilius (2004). “Cell volume response to hyposmotic shock and elevated cAMP in bovine trabecular meshwork cells.” Exp Eye Res 78(1): 15–26.

Stamer, W. D. and A. F. Clark (2017). “The many faces of the trabecular meshwork cell.” Exp Eye Res 158: 112–123.

Stokum, J. A., M. S. Kwon, S. K. Woo, O. Tsymbalyuk, R. Vennekens, V. Gerzanich and J. M. Simard (2018). “SUR1-TRPM4 and AQP4 form a heteromultimeric complex that amplifies ion/water osmotic coupling and drives astrocyte swelling.” Glia 66(1): 108–125.

Sucha, P., Z. Hermanova, M. Chmelova, D. Kirdajova, S. Camacho Garcia, V. Marchetti, I. Vorisek, J. Tureckova, E. Shany, D. Jirak, M. Anderova and L. Vargova (2022). “The absence of AQP4/TRPV4 complex substantially reduces acute cytotoxic edema following ischemic injury.” Front Cell Neurosci 16: 1054919.

Susanna, R., Jr., C. Clement, I. Goldberg and M. Hatanaka (2017). “Applications of the water drinking test in glaucoma management.” Clin Exp Ophthalmol 45(6): 625–631.

Toft-Bertelsen, T. L., D. Krizaj and N. MacAulay (2017). “When size matters: transient receptor potential vanilloid 4 channel as a volume-sensor rather than an osmo-sensor.” J Physiol 595(11): 3287–3302.

Toft-Bertelsen, T. L., O. Yarishkin, S. Redmon, T. T. T. Phuong, D. Krizaj and N. MacAulay (2019). “Volume sensing in the transient receptor potential vanilloid 4 ion channel is cell type-specific and mediated by an N-terminal volume-sensing domain.” J Biol Chem 294(48): 18421–18434.

Tureckova, J., Z. Hermanova, V. Marchetti and M. Anderova (2023). “Astrocytic TRPV4 Channels and Their Role in Brain Ischemia.” Int J Mol Sci 24(8).

Uchida, T., S. Shimizu, R. Yamagishi, S. M. Tokuoka, Y. Kita, R. Sakata, M. Honjo and M. Aihara (2021). “TRPV4 is activated by mechanical stimulation to induce prostaglandins release in trabecular meshwork, lowering intraocular pressure.” PLoS One 16(10): e0258911.

Vriens, J., H. Watanabe, A. Janssens, G. Droogmans, T. Voets and B. Nilius (2004). “Cell swelling, heat, and chemical agonists use distinct pathways for the activation of the cation channel TRPV4.” Proc Natl Acad Sci U S A 101(1): 396–401.

Watanabe, H., J. Vriens, J. Prenen, G. Droogmans, T. Voets and B. Nilius (2003). “Anandamide and arachidonic acid use epoxyeicosatrienoic acids to activate TRPV4 channels.” Nature 424(6947): 434–438.

Yan, X., M. Li, J. Wang, H. Zhang, X. Zhou and Z. Chen (2022). “Morphology of the Trabecular Meshwork and Schlemm’s Canal in Posner-Schlossman Syndrome.” Invest Ophthalmol Vis Sci 63(1): 1.

Yang, G., M. Jia, G. Li, Y. Y. Zang, Y. Y. Chen, Y. Y. Wang, S. Y. Zhan, S. X. Peng, G. Wan, W. Li, J. J. Yang and Y. S. Shi (2024). “TMEM63B channel is the osmosensor required for thirst drive of interoceptive neurons.” Cell Discov 10(1): 1.

Yarishkin, O., T. T. T. Phuong, J. M. Baumann, M. L. De Ieso, F. Vazquez-Chona, C. N. Rudzitis, C. Sundberg, M. Lakk, W. D. Stamer and D. Krizaj (2021). “Piezo1 channels mediate trabecular meshwork mechanotransduction and promote aqueous fluid outflow.” J Physiol 599(2): 571–592.

Yarishkin, O., T. T. T. Phuong, C. A. Bretz, K. W. Olsen, J. M. Baumann, M. Lakk, A. Crandall, C. Heurteaux, M. E. Hartnett and D. Krizaj (2018). “TREK-1 channels regulate pressure sensitivity and calcium signaling in trabecular meshwork cells.” J Gen Physiol 150(12): 1660–1675.

Yarishkin, O., T. T. T. Phuong, F. Vazquez-Chona, J. Bertrand, J. van Battenburg-Sherwood, S. N. Redmon, C. N. Rudzitis, M. Lakk, J. M. Baumann, M. Freichel, E. M. Hwang, D. Overby and D. Krizaj (2022). “Emergent Temporal Signaling in Human Trabecular Meshwork Cells: Role of TRPV4-TRPM4 Interactions.” Front Immunol 13: 805076.

Zeuthen, T. and N. MacAulay (2002). “Passive water transport in biological pores.” Int Rev Cytol 215: 203–230.

Zhang, Y., F. Gao, V. L. Popov, J. W. Wen and O. P. Hamill (2000). “Mechanically gated channel activity in cytoskeleton-deficient plasma membrane blebs and vesicles from Xenopus oocytes.” J Physiol 523 Pt 1(Pt 1): 117–130.

Zhang, Y., P. Liang, L. Yang, K. Z. Shan, L. Feng, Y. Chen, W. Liedtke, C. B. Coyne and H. Yang (2022). “Functional coupling between TRPV4 channel and TMEM16F modulates human trophoblast fusion.” Elife 11.

